# Altered functional brain dynamics in chromosome 22q11.2 deletion syndrome during facial affect processing

**DOI:** 10.1101/2020.12.17.423342

**Authors:** Eli J. Cornblath, Arun Mahadevan, Xiaosong He, Kosha Ruparel, David M. Lydon-Staley, Tyler M. Moore, Ruben C. Gur, Elaine H. Zackai, Beverly Emanuel, Donna M. McDonald-McGinn, Daniel H. Wolf, Theodore D. Satterthwaite, David R. Roalf, Raquel E. Gur, Danielle S. Bassett

## Abstract

Chromosome 22q11.2 deletion syndrome (22q11.2DS) is a multisystem disorder associated with multiple congenital anomalies, variable medical features, and neurodevelopmental differences resulting in diverse psychiatric phenotypes, including marked deficits in facial memory and social cognition. Neuroimaging in individuals with 22q11.2DS has revealed differences relative to matched controls in BOLD fMRI activation during facial affect processing tasks, but time-varying interactions between brain areas during facial affect processing have not yet been studied in 22q11.2DS. We applied constrained principal component analysis to identify temporally overlapping brain activation patterns from BOLD fMRI data acquired during an emotion identification task from 58 individuals with 22q11.2DS and 58 age-, race-, and sex-matched healthy controls. Delayed frontal-motor feedback signals were diminished in individuals with 22q11.2DS, as were delayed emotional memory signals engaging amygdala, hippocampus, and entorhinal cortex. Early task-related engagement of motor and visual cortices and salience-related insular activation were relatively preserved in 22q11.2DS. Insular activation was associated with task performance within the 22q11.2DS sample. Differences in cortical surface area, but not cortical thickness, showed spatial alignment with an activation pattern associated with face processing. These findings suggest that relative to matched controls, primary visual processing and insular function are relatively intact in individuals with 22q11.22DS, while motor feedback, face processing, and emotional memory processes are more affected. Such insights may help inform potential interventional targets and enhance the specificity of neuroimaging indices of cognitive dysfunction in 22q11.2DS.

## INTRODUCTION

Chromosome 22q11.2 deletion syndrome (22q11.2DS) is a genetic neurodevelopmental disorder characterized by a submicroscopic deletion of the long arm of chromosome 22q^1^, which causes a heterogeneous mix of cardiac, endocrine, palatal, immune, gastrointestinal, genitourinary, skeletal, and psychiatric abnormalities^1^. 22q11.2DS is one of the strongest genetic risk factors for psychosis, with over 25% prevalence of psychosis-spectrum symptoms in affected adults^2,3^, alongside co-morbid autism spectrum, attention-deficit, anxiety, and mood symptoms^3,4^. In-depth cognitive phenotyping of individuals with 22q11.2DS suggests that deficits in face memory, affective processing, and social cognition stand out against a backdrop of global cognitive dysfunction^5^.

Facial affect processing relies on the coordination of visual and emotion processing, top-down and bottom-up attention, and memory encoding and retrieval^6,7^. These cognitive processes are subserved by temporally coordinated, evoked activity within a distributed network of limbic, insular, visual, and medial and lateral prefrontal brain areas^6,8,9^. Neuroimaging studies have implicated early top-down inhibition from anterior cingulate cortex to the amygdala in facial affect processing^9,10^. Increased amygdalar activation to stimuli with negative emotional salience has been found in schizophrenia^11,12^, along with reduced interaction between amygdala and frontoparietal cortex^13^. Nevertheless, it remains unclear how regional activations and network interactions result in behaviorally relevant emotion processing, and which components of this process contribute to dysfunctional facial affect processing in 22q11.2DS.

Multi-modal neuroimaging phenotypes in 22qll.2DS have not provided clear explanations for the observed behavioral abnormalities or identified candidates for targeted intervention^1,3^. T1-weighted structural imaging has revealed altered cortical thickness and surface area, with the strongest differences affecting midline, lateral inferior frontal, and superior parietal cortex^14^. Resting state fMRI (rs-fMRI) studies have found differences in default mode network^15,16^ and frontolimbic connectivity, the latter of which correlates with anxiety^17^, suggesting that frontolimbic dysconnectivity is relevant to affect processing in 22q11.2DS. Task-based fMRI studies of facial affect processing in 22q11.2DS have revealed reduced amygdalar fear accommodation and fusiform gyrus activation^18,19^; however, these studies are limited by their focus on univariate activation measures, given that facial affect processing inherently relies on interactions among brain regions.

Here, we hypothesized that primary visual and motor processing would be preserved in individuals with 22q11.2DS, while frontolimbic interactions subserving bottom-up emotion-processing^6,7^ would be disrupted in individuals with 22q11.2DS. We applied constrained principal component analysis (CPCA)^13,20–24^ to identify brain activation patterns evoked by images of faces, and quantified their time course of activation after emotion identification. Specifically, we used emotion identification task fMRI data^8,11,25^ acquired from 58 individuals with 22q11.2DS identified through the 22q and You Center at the Children’s Hospital of Philadelphia, examined as part of a prospective brain-behavior study of 22q11.2DS, and 58 age-, sex-, and race-matched healthy controls (HCs) from the Philadelphia Neurodevelopmental Cohort^26,27^. The spatial profiles of these activation patterns were similar between groups, but their temporal profiles were altered in 22q11.2DS, implicating selective dysfunction in putative motor feedback (PC2) and emotional memory (PC5) signals. PC2 and PC4 activation were most strongly associated with task performance within the 22q11.2DS sample. Finally, we quantified the alignment between these task-evoked spatial activation patterns and spatial maps of gray matter structural change in individuals with 22q11.2DS. Collectively, these findings shed light on the dynamic interactions between visual, attentional, limbic, and motor systems during facial affect processing and distinguish between affected and relatively unaffected task-relevant neural systems in individuals with 22q11.2DS.

## METHODS

### Participants

Emotion identification task fMRI data were obtained from a sample of 58 individuals with genotype-confirmed chromosome 22q11.2 deletion syndrome evaluated by the 22q and You Center at the Children’s Hospital of Philadelphia and the Philadelphia Neurodevelopmental Cohort (PNC)^26^, a large community-based study of brain development (see Table I). Here, we study a sample of *n* = 58 age-, sex-, and race-matched PNC subjects without radiological abnormalities or medical problems that might impact brain function. All subjects in this sample had a mean framewise displacement < 0.5 mm during the emotion identification task to minimize motion-related confounds.

**TABLE I.** Sample characteristics. The *p*-value column was generated using two independent sample *t*-tests, except for proportions of race, which were generated by comparing bootstrapped confidence intervals of sample proportions of each race. All values, except race and sex, are represented as a mean ± standard deviation. *CNB*, mean *z*-scored accuracy across all Penn Computerized Neurocognitive Battery sections as a surrogate for intelligence quotient^28^. *NR*, no response.

|  | 22q11.2DS | PNC | p-value |
| --- | --- | --- | --- |
| *Demographics* |  |  |  |
| Age (y) | 20.3 ± 4.8 | 19.6 ± 3.9 | 0.38 |
| Male | 50% | 50% | - |
| White | 81% | 75.9% | 0.58 |
| African American | 12.1% | 17.2% | 0.51 |
| Other Race | 6.9% | 6.9% | 1 |
| CNB Accuracy (z) | -1.2 | 0.22 | $8.3 \times 10^{-20}$ |
| Typical or atypical antipsychotics, $n$ (%) | 5 (8.6%) | - | - |
| *Imaging* |  |  |  |
| Mean Framewise Displacement (mm) | 0.119 ± 0.077 | 0.0762 ± 0.085 | 0.0057 |
| Total Brain Volume (cm<sup>3</sup>) | 1110 ± 120 | 1220 ± 120 | 2.3e-06 |
| *Task Performance (%)* |  |  |  |
| Correct | 72.9 ± 21 | 90.9 ± 6.2 | $4.4 \times 10^{-8}$ |
| Incorrect | 17.4 ± 15 | 6.73 ± 4.6 | $3.9 \times 10^{-6}$ |
| NR | 7.92 ± 13 | 2.31 ± 4.3 | 0.0033 |
| Threat Correct | 70.1 ± 23 | 89.1 ± 10 | $2.8 \times 10^{-7}$ |
| Threat Incorrect | 19.6 ± 18 | 8.33 ± 7.6 | $4.7 \times 10^{-5}$ |
| Threat NR | 8.48 ± 13 | 2.56 ± 5.7 | 0.0029 |
| Non-Threat Correct | 74.8 ± 21 | 92.1 ± 5 | $1.5 \times 10^{-7}$ |
| Non-Threat Incorrect | 15.9 ± 15 | 5.66 ± 3.9 | $4.7 \times 10^{-6}$ |
| Non-Threat NR | 7.54 ± 14 | 2.14 ± 4.1 | 0.007 |

### Emotion identification task

As previously described^8,11,25^, the emotion identification task employed a fast event-related design with a jittered inter-stimulus interval (ISI). Subjects viewed 60 faces displaying neutral, happy, sad, angry, or fearful expressions, and were asked to label the emotion displayed. Stimuli construction and validation are detailed elsewhere^29^. Briefly, the stimuli were color photographs of actors (50% female) who volunteered to participate in a study on emotion. They were coached by professional directors to express a range of facial expressions. For the present task, a subset of intense expressions was selected based on high degree of accurate identification (80%) by raters. Each face was displayed for 5.5 seconds followed by a variable ISI of 0.5 to 18.5 seconds, during which a crosshair (matching the faces’ perceptual qualities) was displayed. Total task duration was 10.5 minutes.

### Structural and functional image processing

We used *fMRIprep* software^30^ to perform brain extraction and segmentation of the individual high-resolution T1-weighted images, registration of task fMRI BOLD volumes to individual-specific T1 images, and computation of confound time series (see Supplementary Information for *fMRIprep* standardized methods section). After the above steps were completed using *fMRIprep* software^30^, we used XCP engine^31^ to perform the following steps: (1) demeaning to remove linear or quadratic trends, (2) first-order Butterworth filtering to retain signal in the 0.01 to 0.50 Hz range, and (3) confound regression of 6 realignment parameters. Following these preprocessing steps, we extracted parcellated, regional time series from the unsmoothed voxel-level data using the 200-node Schaefer cortical atlas^32^ and 14 subcortical nodes defined by the Harvard-Oxford atlas^33^.

### Extracting task-relevant spatiotemporal modes of brain activity through constrained principal component analysis

After completing the outlined preprocessing steps, we used constrained principal components analysis (CPCA)^23,24^ to extract task-evoked spatial modes of brain activation at the group-level with subject-level temporal weights^13,20–22^. Briefly, this approach involves using a finite impulse response (FIR) basis set^34^ to extract task-related variance from a set of BOLD timeseries, applying principal component analysis (PCA) to extract orthogonal spatiotemporal modes from the task-related variance, and then a second regression step using the same FIR basis set to determine how the temporal scores of each PC fluctuate with stimulus presentation. Here, our FIR basis set contained an indicator variable for each image acquisition spanning 3-18 seconds after each of 6 task events, consisting of correct, incorrect, and non-responses to threatening and non-threatening stimuli. See Supplementary Information for additional details and mathematical formulation.

### Multilevel growth models of principal component response curves

In order to compare the activation of each CPCA component evoked by each task event between HCs and individuals with 22q11.2DS, we applied a multilevel growth modeling approach. This approach allowed us to account for the multilevel nature of the data, with multiple time points of component activity for different stimuli nested within participants, as well as between-subject factors such as age and sex. Briefly, for each component, we used the `nlme` package in R to fit a linear mixed effects model predicting the estimated score of that component at time *t* after each task event, excluding the 2 non-response task events. All models included age, sex, total brain volume, mean framewise displacement during task scans, handedness, and group membership (22q11.2DS or control). All models additionally included random intercepts to capture between-person differences in mean levels of component scores.

Next, a partially supervised model selection procedure motivated by a previous study^35^ was implemented in order to include fixed effects of time, polynomials of time, stimulus type, response type, and interactions between those variables and 22q11.2DS status, when the inclusion of these variables improved model fit. We also included random effects of time to model between-person differences in how activity changed across time. See Supplementary Information, “Multilevel growth model selection procedure” for more details.

## RESULTS

### Identifying brain activation patterns evoked by emotion identification

Individuals with 22q11.2DS exhibit deficits in facial affect processing and social cognitive function. However, the dynamic patterns of brain activation underlying these deficits are not fully understood. Here, we conducted a spatiotemporally sensitive analysis of task-related brain activity using constrained principal component analysis (CPCA)^13,20,21^ to analyze BOLD data from 58 individuals with 22q11.2DS and 58 age-, sex-, and race-matched healthy controls (HCs). First, we regressed BOLD signal (Fig. 1a) from an emotion identification task onto a finite impulse response (FIR) basis set to extract stimulus-related signals (Fig. 1b). We used separate regressors for each subject and 4 *task events* of interest, in which threatening or non-threatening stimuli were accompanied by either correct or incorrect responses^36,37^. Next, to complete the CPCA procedure, we identified the principal components of the task related variance in BOLD signal captured by the predicted values of this regression (Fig. 1c). A scree plot of the variance explained by this principal component analysis revealed an elbow at 6 components, which cumulatively explained 64.1% of the task-related variance in the BOLD signal (Fig. S1a). The first principal component (Fig. S2), explaining 36.7% of task-related variance, appeared to reflect a global signal fluctuation^38^, and was thus excluded from further analysis. We named this global signal component “PC0” and re-indexed the original PC2-6 as PC1-5 for future analyses. Finally, we applied a bootstrapping analysis (see Supplementary Information, subsection “Bootstrapping analysis of CPCA components”) to threshold these spatial maps (Fig. 2a) and demonstrate that a group CPCA solution was adequate to describe each cohort’s BOLD data (Fig. S3a). Collectively, these analyses revealed multiple task-evoked spatial activity patterns that occur in both HCs and individuals with 22q11.2DS.

**FIG. 1.**
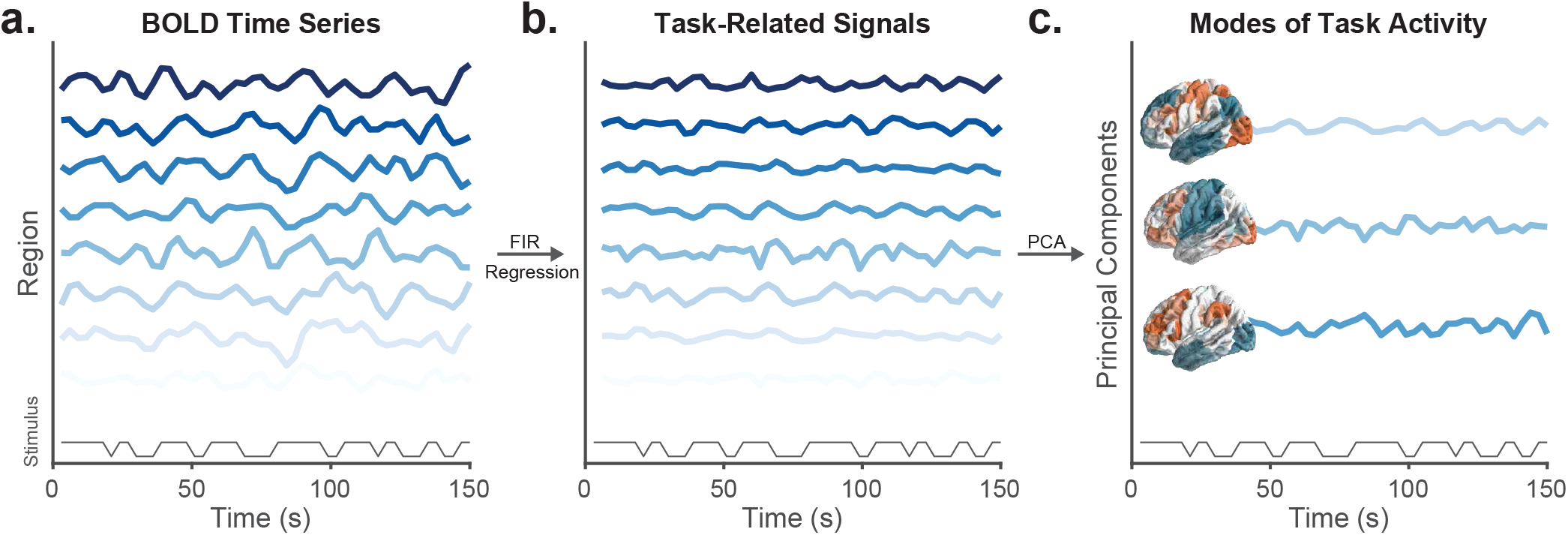
Schematic of methods for functional image analysis. *(a)* Example time series of BOLD signal from 7 arbitrarily chosen regions acquired during an emotion identification task. Boxcar regressor of stimulus presentation is shown below the BOLD signal. *(b)* In order to isolate task-related signals, the BOLD signal from panel *a* is regressed onto a finite impulse response basis set, which flexibly captures each region’s response to different stimuli without assuming any particular shape of the hemodynamic response function. *(c)* The predicted values of the linear regression model are decomposed with principal component analysis, yielding orthogonal spatial maps of task-evoked brain activity with orthogonal temporal profiles. These spatiotemporal modes can be related back to stimulus presentation in order to estimate the task evoked time course of each spatial activation pattern. *FIR*, finite impulse response. *PCA*, principal component analysis.

**FIG. 2.**
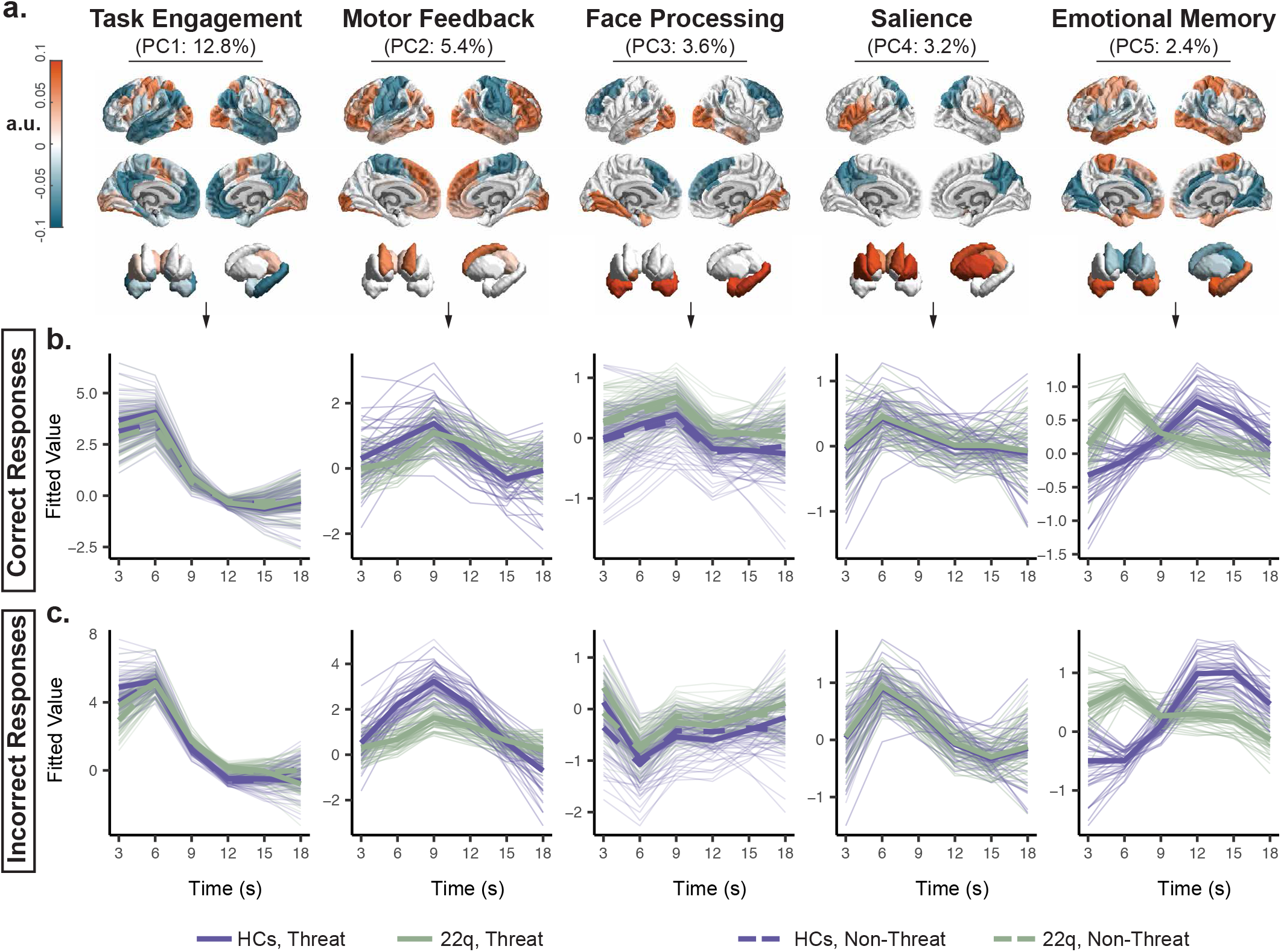
Spatiotemporal modes of activity evoked by emotion identification are selectively altered in 22q11.2DS. *(a)* Spatial loadings of the first 5 principal components of task-related variance (Fig. 1b) in emotion identification task BOLD signal thresholded at *p* < 10^−4^ using bootstrap significance testing^41^, shown on surface renderings of cortex and subcortex. Components are named for the cognitive process they putatively reflect. *(b, c)* Multilevel growth models fit to the temporal scores (*y*-axis) of each task-evoked PC during the time (*x*-axis) occurring 3-18 seconds after correct (panel *b*) or incorrect (panel *c*) emotion identification of threatening (thick lines) and non-threatening (dashed lines) faces. We used a model selection procedure (see Methods) to predict each PC’s scores over time from polynomials of time, stimulus type (threat or non-threat), response type (correct or incorrect), 22q status, and interactions between those variables while controlling for age, sex, total brain volume, head motion, and handedness. The best model selected through this process was used to obtain fitted values (*y*-axis) to describe the trajectory of each PC’s score for the prototypical individual in each group (thick, opaque lines) and for each participant (thin, faded lines).

### Altered temporal profiles of task-general brain activity in 22q11.2DS

After identifying spatial patterns of task-related brain activity, we next sought to characterize each signal component’s evoked response to the 4 *task events*. We regressed PC scores onto an FIR basis set to estimate the mean score of each PC at the 6 image acquisitions occurring 3-18 seconds after each task event (Fig. S4a,c). Next, we applied a model selection procedure using multilevel growth models to parameterize the shape of each PC’s event response curve with polynomial functions of time (Fig. 2b,c; see Methods). This analysis allowed us to statistically compare the temporal profiles of these PC response curves between HCs and 22q11.2DS individuals while accounting for effects and interactions (Supplementary Data File 1) of between-subject factors (total brain volume, sex, age, head motion, and handedness) and within-subject factors (task event). Notably, results were robust to parcellation scheme (Fig. S5) and no activation was detected when BOLD data were phase randomized to create stimulus-independent surrogate null data (Fig. S6).

First, we observed that PC1 was rapidly and robustly engaged in each task event, peaking around 6 seconds after task event onset (Fig. 2c,e, leftmost subpanel). The spatial map of PC1 revealed default mode network deactivation^39,40^, visual cortex activation, and left-hemispheric hand motor cortex activation. The temporal expression of PC1 was highest during correct responses to threat stimuli (Fig. 2b,c; Time^3^ × Threat × Correct, *β* = 4.8 × 10^−3^, *p* = 3.7 × 10^−3^, *df* = 2200), but primarily differed between HCs and 22q11.2DS during incorrect responses and less so during correct responses (Fig. 2b,c; Time^2^ ×22q×Correct, *β* = 0.021, *p* =1.5 × 10^−3^, *df* = 2200).

Next, we observed that PC2 showed the most pronounced activation during incorrect responses (Fig. 2b,c; Time^2^×Correct, *β* = 0.037, *p* = 5.2 × 10^−16^, *df* = 2200). The PC2 peak was delayed, occurring around 9 seconds after the task event in contrast to the peak at 6 seconds observed in PC1. The spatial map of PC2 consisted of dorsolateral and ventrolateral prefrontal cortex activation amid low amplitude activity in sensorimotor areas. Notably, we found an interaction between 22q11.2DS status, time, and response type such that 22q11.2DS showed reduced activation of PC2 during incorrect responses (Fig. 2b,c; Time^2^×22q×Correct, *β* = −0.031, *p* =6.3 × 10^−7^, *df* = 2200).

PC3 activity showed a positive peak around 9 seconds during correct responses and a negative peak at 6 seconds during incorrect responses, with the greatest responses to threatening stimuli (Fig. 2b,c; Time^2^ ×Correct×Threat, *β* = −0.012, *p* = 0.031, *df* = 2200). The spatial map of PC3 showed activation of the amygdala, hippocampus, and fusiform gyrus, with activity decreases in dorsolateral prefrontal regions (Fig. 2a). In 22q11.2DS, activation of this component was higher at baseline (Fig. 2b,c; 22q, *β* = 0.29, *p* = 1.2 × 10^−3^, *df* = 96), apparently capturing the attenuated decrease of this component during incorrect response (Fig. S4c, third panel from right).

PC4 peaked early around 6 seconds after the task event. The spatial map of PC4 was characterized by activation in the bilateral opercula, insulae, and motor basal ganglia with low amplitude activity in the posterior cingulate and posterior parietal cortex (Fig. 2a). Stimulus type was not associated with PC4’s time course, but the response was more pronounced during incorrect trials. (Fig. 2b,c; Time^3^ × Correct, *β* = −2.2 × 10^−3^, *p* = 5 × 10^−4^, *df* = 2200). There was a trend towards reduced PC4 expression during correct non-threat trials in HCs only (Fig. S4a, 4^th^ panel from the left), but models containing time-by-stimulus-by-response-by-cohort interaction coefficients did not meet statistical significance. Overall, we did not detect any statistically significant group differences in the temporal response of PC4.

Finally, PC5 exhibited the most delayed activation profile, peaking around 12 seconds after the task event in HCs (Fig. 2b,c; Time^5^, *β* = 8 × 10^−5^, *p* = 0.012, *df* = 2200). However, this peak was shifted to 6 seconds after the task event in individuals with 22q11.2DS (Fig. 2b,c; Time^4^×22q, *β* = −5.5 × 10^−4^, *p* = 5.4 × 10^−4^, *df* = 2200). The spatial map of PC5 showed engagement of the hippocampus, amygdala, entorhinal cortex, ventromedial prefrontal cortex, and bilateral hand motor sensorimotor cortices (Fig. 2a).

### Individual differences in activation peaks explain variance in task performance within 22q11.2DS sample

Next, we were interested to understand the relevance of these spatiotemporal modes of brain activation to cognitive function within the 22q11.2DS population. We used each 22q11.2DS individual’s peak score on each of the 5 components during each of the 4 task events as independent variables in separate models to predict the rank of in-scanner accuracy on the emotion identification task under study. We used the rank of accuracy as our outcome variable rather than the percentage accuracy in order to include as many 22q11.2DS individuals as possible without biasing regression estimation with outlier values. Each of these 20 models included age, sex, total brain volume, mean task-scan head motion, and handedness as covariates.

This analysis revealed that PC2 and PC4 scores were the most strongly associated with correct emotion identification in 22q11.2DS individuals. Specifically, we found that PC2 peak values during threat incorrect (Fig. 3a; *β* = 0.55, *p*_FDR_ = 0.0022, *df* = 42) and non-threat incorrect trials (Fig. 3a, b; *β* = 0.63, *p*_FDR_ = 1.99 × 10^−5^, *df* = 43) were positively associated with emotion identification accuracy. PC4 peak values during non-threat correct trials were negatively associated with accuracy (Fig. 3a,c; *β* = −0.45, *p*_FDR_ = 0.0075, *df* = 45), whereas PC4 peak values during non-threat incorrect trials were positively associated with accuracy (Fig. 3a; *β* = 0.49, *p*_FDR_ = 0.0055, *df* = 43). These findings suggest that the presence of putative motor feedback signals (Fig. 2) during incorrect trials and apparent salience-related insular activation (Fig. 2) during incorrect trials but not correct trials index accurate emotion identification in 22q11.2DS.

**FIG. 3.**
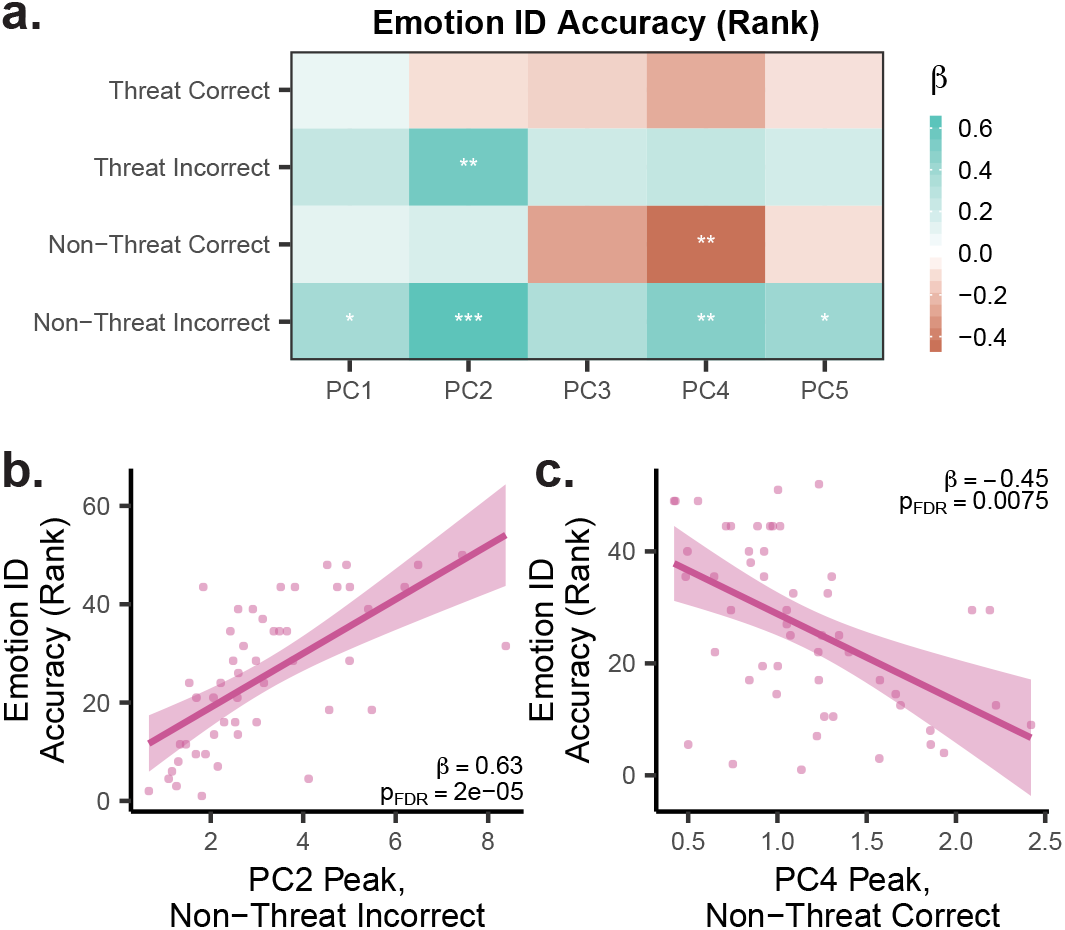
Overall task performance in individuals with 22q11.2DS can be predicted from peak PC scores. *(a)* Standardized linear regression *β* weights (color axis) for the peak value of each PC (*x*-axis) during each task event (*y*-axis) as a predictor of overall in-scanner emotion identification accuracy using the sample of individuals with 22q11.2DS only, in a model containing age, sex, total brain volume, head motion, and handedness as covariates. Asterisks indicate level of significance after FDR correction (*q* < 0.05) over all 20 *β* values: *, *p*_FDR_ < 0.05. **, *p*_FDR_ < 0.01. ***, *p*_FDR_ < 0.001. *(b-c)* Partial residuals of emotion identification accuracy (*y*-axis) from linear regression models in panel *a* plotted against peak PC2 scores during incorrect responses to non-threatening stimuli (panel *b*) or peak PC4 scores during correct responses to non-threatening stimuli (panel *c*) (*x*-axis).

### Differences in brain structure in 22q11.2DS selectively align with task-evoked activation patterns

After characterizing functional brain abnormalities during emotion identification in 22q11.2DS, we examined whether differences in gray matter morphometry could be a substrate for these functional effects. Here, we tested the hypothesis that areas with abnormal cortical morphometry in 22q11.2DS align with the identified task-evoked activation patterns, possibly hindering the function of regions that are specifically engaged during emotion identification (Fig. 2b-e).

To test this hypothesis, we utilized difference maps of cortical thickness (Fig. 4a) and cortical surface area (Fig. 4b) obtained from a previously published manuscript^14^ using a larger, partially overlapping sample. We computed the mean absolute value (MAV) of structural change for each metric within the cortical areas of each spatial PC map for which the loading value was significantly different from 0 after bootstrap thresholding (Fig. 2a, *p* < 10^−4^). This metric captures the total extent of structural differences within activated or deactivated regions for each PC. We compared the MAV values (Fig. 4a-b, yellow diamonds) to a null distribution of MAV values obtained using 500 permuted structural maps with preserved spatial covariance^42^. This analysis revealed that PC3 harbored differences in cortical surface area within its engaged areas that were greater than expected due to spatial covariance alone (Fig. 4b; MAV= 0.059, *p*_spin_FDR__ < 0.002). The MAV of cortical thickness within any PC map did not differ from that which would be expected due to spatial covariance (Fig. 4a; all *p*_spin_FDR__ > 0.05). These findings suggest that differences in cortical surface area, rather than cortical thickness, align more specifically with activation patterns associated with face processing.

**FIG. 4.**
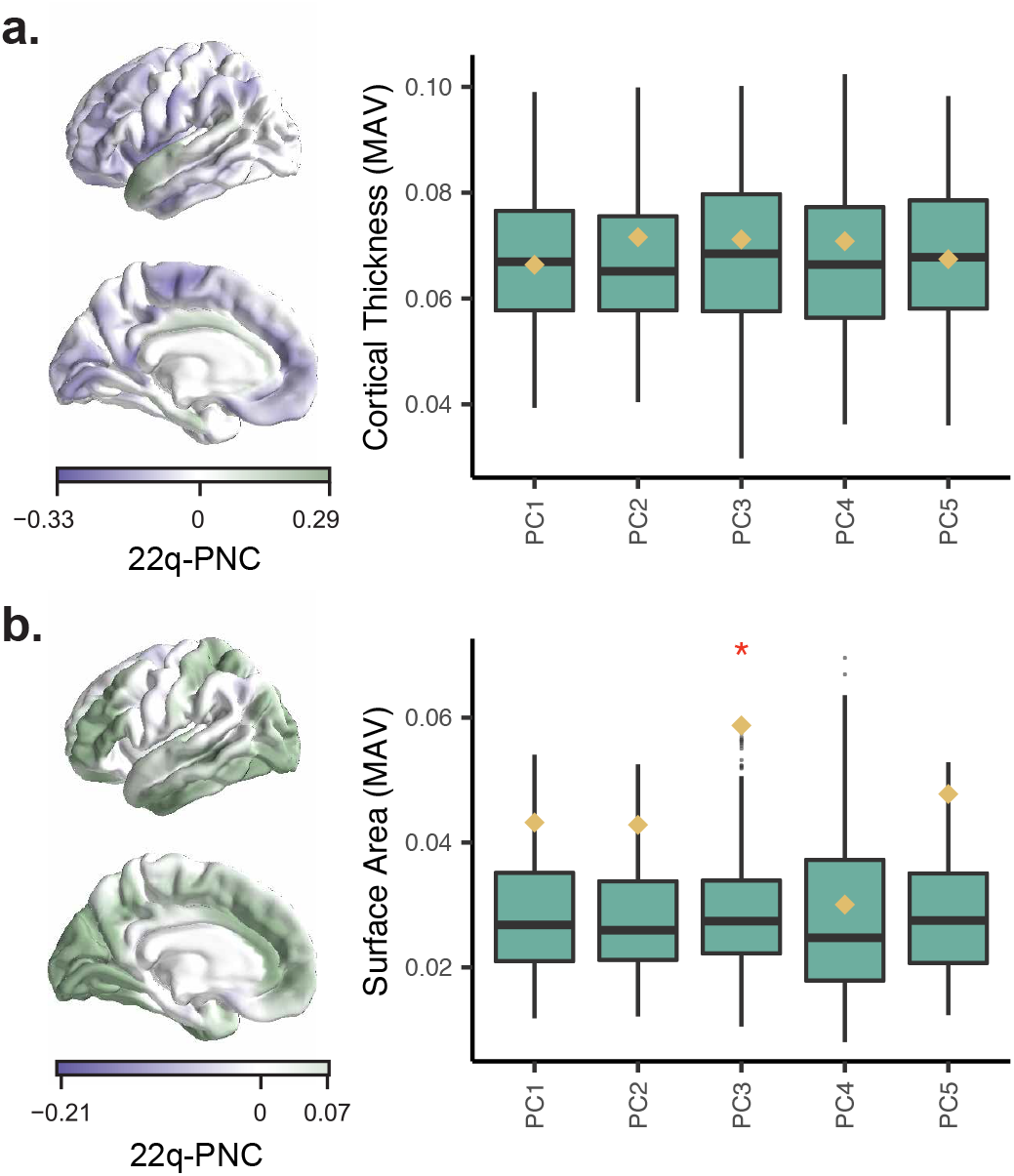
Differences in cortical surface area in 22q11.2DS align with face processing component. *(a-b)* Surface plots show cortical thickness (panel *a*) and cortical surface area (panel *b*) differences between HCs and individuals with 22q11.2DS, reproduced with permission from Sun *et al*. 2018^14^. Yellow diamonds show mean absolute value (MAV) of cortical thickness (panel *a*) and cortical surface area (panel *b*) differences within the areas of each spatial PC map (Fig. 2a) that differed from 0 after bootstrap thresholding at *p* < 10^−4^. The green boxplots show the same measure of MAV within each PC map computed using 500 permuted versions of the structural maps with preserved spatial covariance^42^. Red *, *p*_spin_FDR__ < 0.05, corrected over 12 comparisons for 6 PCs and 2 structural maps.

## DISCUSSION

In the present study, we extracted 5 spatial patterns of task-evoked brain activity from individuals with 22q11.2DS and matched healthy controls. These activation patterns appeared to engage both “task-general” (PC1, PC2, and PC4) systems that are seen across many tasks, as well as “emotion-related” (PC3 and PC5) systems, which are more specifically engaged during facial affect processing tasks. We found the strongest group differences in PC2 and PC5. Finally, we showed cortical gray matter surface area differences in 22q11.2DS aligned with the spatial map of PC3, due to engagement of primary visual cortex, inferotemporal cortex, and dorsolateral prefrontal cortex.

### Altered task-general brain dynamics in 22q11.2DS

Of the three task-general components, we found that PC1 and PC4 were relatively preserved in 22q11.2DS. PC1 contained rapid engagement of dominant hand motor cortex with visual cortex activation and default mode deactivation observed across all task events. Default mode (DM) deactivation is a hallmark of goal oriented tasks^40^. Prior fMRI studies in 22q11.2DS have found both decreased and increased spontaneous activity in DM subregions^16,43^. In the task fMRI setting studied here, we find timing dependent differences. The DM is relatively unaffected in the early response (PC1) with more group differences in DM subregions in the delayed response (PC2 and PC5). PC4 was characterized by early-peaking insular activation, most robust after incorrect responses to potentially unfamiliar or ambiguous stimuli, consistent with the insula’s role in detecting novel stimuli^44^. Emotion identification accuracy was negatively associated with PC4 activation during incorrect trials within the 22q11.2DS sample, suggesting that inappropriate, early insular responses to stimuli may contribute to or reflect poor task performance.

The remaining task-general component implicates aberrant motor feedback in 22q11.2DS during incorrect responses to failures of emotion identification. In HCs, PC2 was more strongly engaged during incorrect responses and consisted of delayed frontal activation with sensorimotor deactivation, which we interpreted as a negative feedback signal from bilateral inferior frontal gyri to motor cortex. This pattern is consistent with the known role of the inferior frontal gyrus in response inhibition^45^. The lack of this signal was associated with poor emotion identification accuracy in the 22q11.2DS sample, consistent with previously observed motor dysfunction in individuals with 22q11.2DS^46–48^. However, given that subjects are not notified of incorrect responses during this task, PC2 activation may also be explained by a lack of post-response recognition of an incorrect choice in 22q11.2DS individuals.

### Altered emotion-related brain dynamics in 22q11.2DS

Individuals with 22q11.2DS show deficits in social cognition and face memory even after adjusting for global cognitive deficits^5^. fMRI studies of face processing in 22q11.2DS have found hypoactivation of fusiform gyrus^18,19^ and a lack of amygdalar fear accommodation^18^. Here, we found altered time courses of PC3 and PC5, which both engaged fusiform gyrus, amygdala, and hippocampus. PC3 also revealed dorsolateral prefrontal cortex deactivation and peaked at 9 seconds during correct responses. Interestingly, a negative PC3 peak occurred 6 seconds after incorrect responses, implicating suppression of face processing circuitry and activation of dorsal attention areas in incorrect responses, an effect that was less pronounced in 22q11.2DS. This finding may reflect incorrect responses in HCs resulting from futile goal-directed cognition amidst failure of limbic processing, while individuals with 22q11.2DS may experience failure of limbic processing with less compensatory goal-directed cognition. Additionally, the spatial map of PC3 showed statistically significant alignment with cortical surface area alterations in 22q11.2DS, which may explain the abnormal temporal profile of PC3; however, the observed differences were small, and therefore structural alterations may instead alter local processing despite relatively normal onset. Abnormal local processing could in turn affect the engagement of concurrently (PC2) or later peaking (PC5) components.

In addition to the primary sensory processing underlying facial recognition, emotional memory^49^ contributes to facial affect processing and engages a similar set of brain areas^7^. PC5 harbored thalamic deactivation and a delayed peak around 12-15 seconds, suggesting that this component may reflect memory encoding rather than retrieval in HCs; however, in individuals with 22q11.2DS, this component peaked early at 6 seconds with an absent late peak. This early peak may reflect inappropriate early activation of emotional circuitry and the absence of a late peak may reflect dysfunctional emotional memory encoding. Indeed, emotional memory deficits in a mouse model of 22q11.2DS have been linked to disrupted thalamo-amygdalar signaling^50^. Collectively, PC3 and PC5 may provide separable measures of dysfunction in affective processing in individuals with 22q11.2DS.

### Methodological limitations

Though this study provides a great deal of information about spatiotemporal patterns of task-evoked brain activity in 22q11.2DS, several key limitations must be acknowledged. First, the fact that global cognitive deficits are observed in individuals with 22q11.2DS raises the possibility that reduced task engagement may confound our observations of abnormal task-related brain activity. While we cannot eliminate this possibility, the relative similarity in PC1 activation between groups suggests that primary visual processing, default mode deactivation, and motor execution are intact in 22q11.2DS. Second, PCA enforces a spatiotemporally orthogonal solution, a constraint that is not biologically necessitated. Future studies could explore this limitation by benchmarking PCA solutions against varimax-rotated PCA, nonnegative matrix factorization, or other non-orthogonal decompositions. Finally, individuals with 22q11.2DS exhibit increased in-scanner head motion, and even our relatively lenient motion exclusion threshold (mean framewise displacement < 0. 5mm vs. 0.2 mm^51^) may have biased our sample towards less severe phenotypes in 22q11.2DS. We attempted to address any remaining motion contamination by including mean framewise displacement as a covariate in subsequent regression analysis.

### Future directions

In the future, targeted task design would enhance the interpretation of these signals in relation to emotional cognition in 22q11.2DS. For instance, one could follow the emotion identification task with a face recognition task^36^. If PC5 scores during emotion identification predicts future correct recognition, one could infer that PC5 reflects memory encoding. This task would also allow separation of components involved in emotion identification from those involved in emotion perception. To investigate the relationship between PC2 and motor feedback, one could test whether notification of errors modifies the response of PC2 during incorrect trials.

In the present study, our comparison of structure and function was limited to gray matter differences, though it has been shown that the dynamic spreading of activation along white matter tracts supports task-related and spontaneous fluctuations in brain activity^52–54^. Network control theory^55–57^ provides tools that account for both external inputs, such as task stimuli^53^, and internal spreading dynamics along white matter connections. One recent study found that control properties of structural brain networks explained dysfunctional resting state connectivity in 22q11.2DS^58^; future studies could apply these tools to assess the temporal alterations in stimulus-driven brain activity identified here.

## CODE AVAILABILITY

All analysis code is available at https://github.com/ejcorn/fir_pca_22q.

## CITATION DIVERSITY STATEMENT

Recent work in several fields of science has identified a bias in citation practices such that papers from women and other minorities are under-cited relative to the number of such papers in the field^59–64^. Here we sought to proactively consider choosing references that reflect the diversity of the field in thought, form of contribution, gender, and other factors. We obtained predicted gender of the first and last author of each reference by using databases that store the probability of a name being carried by a woman^59,65^. By this measure (and excluding self-citations to the first and last authors of our current paper), our references contain 12.1% woman(first)/woman(last), 7.6% man/woman, 21.2% woman/man, and 59.1% man/man. This method is limited in that a) names, pronouns, and social media profiles used to construct the databases may not, in every case, be indicative of gender identity and b) it cannot account for intersex, non-binary, or transgender people. We look forward to future work that could help us to better understand how to support equitable practices in science.

## ACKNOWLEDGEMENTS

D.S.B. and E.J.C. acknowledge support from the John D. and Catherine T. MacArthur Foundation, the Alfred P. Sloan Foundation, the ISI Foundation, the Paul Allen Foundation, the Army Research Laboratory (W911NF-10-2-0022), the Army Research Office (Bassett-W911NF-14-1-0679, Grafton-W911NF-16-1-0474), the National Institute of Mental Health (2-R01-DC-009209-11, R01 - MH112847, R01-MH107235, R21-M MH-106799), the National Institute of Child Health and Human Development (1R01HD086888-01), National Institute of Neurological Disorders and Stroke (R01 NS099348), and the National Science Foundation (NSF PHY-1554488, BCS-1631550, and IIS-1926757). E.J.C. acknowledges support from the National Institute of Mental Health (F30 MH118871-01). T.D.S. acknowledges support from the National Institute of Mental Health (R01MH107703, R01MH113550, and RFMH116920). R.E.G. acknowledges support from the National Institute of Mental Health (U01 087626, U01 101719, U01 119738). D.M.M. acknowledges support from the National Institute of Mental Health (U01-MH191719; MH119737-02; R01-MH087636-01A1). D.M.L. acknowledges support from the National Institute on Drug Abuse (K01DA047417). The authors would like to thank Dr. Carrie Bearden and Dr. Frank Daqiang Sun for providing us with cortical thickness and cortical surface area maps from Sun *et al*. 2018^14^. The content is solely the responsibility of the authors and does not necessarily represent the official views of any of the funding agencies.

## SUPPLEMENTARY INFORMATION

### Image acquisition

MRI data were acquired on a 3 Tesla Siemens Tim Trio whole-body scanner and 32-channel head coil at the Hospital of the University of Pennsylvania. High-resolution T1-weighted images (TR = 1810 ms, TE = 3.51 ms, FOV = 180 × 240 mm, matrix = 256 × 192, 160 slices, TI = 1100 ms, flip angle = 9 degrees, effective voxel resolution of 0.9375 × 0.9375 × 1mm) were acquired for each subject. All subjects underwent functional imaging (TR = 3000 ms; TE = 32 ms; flip angle = 90 degrees; FOV = 192 × 192 mm; matrix = 64 × 64; slices = 46; slice thickness = 3 mm; slice gap = 0 mm; effective voxel resolution = 3.0 × 3.0 × 3.0 mm) during the emotion-identification task sequence^26^. Throughout the study, subjects’ heads were stabilized in the head coil using one foam pad over each ear and a third pad over the top of the head in order to minimize motion. The emotion identification task was displayed using Presentation software, and both responses and response times were recorded using a custom fiberoptic response pad. Prior to any image acquisition, subjects were acclimated to the MRI environment via a mock scanning session in a decommissioned scanner. Mock scanning was accompanied by acoustic recordings of gradient coil noise produced by each scanning pulse sequence. Feedback regarding head motion was provided using the MoTrack motion tracking system (Psychology Software Tools, Inc., Sharpsburg, PA).

### Image processing

Results included in this manuscript come from preprocessing performed using *fMRIPrep* 1.5.8^30,66^ (RRID:SCR_016216), which is based on *Nipype* 1.4.1^67,68^ (RRID:SCR_002502).

#### Anatomical data preprocessing using fMRIprep software

The T1-weighted (T1w) image was corrected for intensity non-uniformity (INU) with `N4BiasFieldCorrection`^69^, distributed with ANTs 2.2.0^70^ (RRID:SCR_004757), and used as the T1w-reference throughout the workflow. The T1w-reference was then skull-stripped with a *Nipype* implementation of the `antsBrainExtraction.sh` workflow (from ANTs), using OASIS30ANTs as target template. Brain tissue segmentation of cerebrospinal fluid (CSF), whitematter (WM) and gray-matter (GM) was performed on the brain-extracted T1w using `fast`^71^. Brain surfaces were reconstructed using `recon-all`^72^, and the brain mask estimated previously was refined with a custom variation of the method to reconcile ANTs-derived and FreeSurfer-derived segmentations of the cortical gray-matter of Mindboggle^73^. Volume-based spatial normalization to one standard space (MNI152NLin2009cAsym) was performed through nonlinear registration with `antsRegistration` (ANTs 2.2.0), using brain-extracted versions of both T1w reference and the T1w template. The following template was selected for spatial normalization: *ICBM 152 Nonlinear Asymmetrical template version 2009c* [^74^, RRID:SCR_008796; TemplateFlow ID: MNI152NLin2009cAsym].

#### Functional data preprocessing using fMRIprep software

For each BOLD run found per subject (across all tasks and sessions), the following preprocessing was performed. First, a reference volume and its skull-stripped version were generated using a custom methodology of *fMRIPrep*. Susceptibility distortion correction (SDC) was omitted. The BOLD reference was then co-registered to the T1w reference using `bbregister` (FreeSurfer) which implements boundary-based registration^75^. Co-registration was configured with six degrees of freedom. Head-motion parameters with respect to the BOLD reference (transformation matrices, and six corresponding rotation and translation parameters) are estimated before any spatiotemporal filtering using MCFLIRT `Jenkinson2002`^76^. BOLD runs were slice-time corrected using `3dTshift` from AFNI 20160207^77^ (RRID:SCR_005927). The BOLD time-series were resampled to surfaces on the following spaces: *fsaverage5*. The BOLD time-series (including slice-timing correction when applied) were resampled into their original, native space by applying the transforms to correct for head-motion. These resampled BOLD time-series will be referred to as *preprocessed BOLD in original space*, or just *preprocessed BOLD*. The BOLD time-series were resampled into standard space, generating a *preprocessed BOLD run in [‘MNI152NLin2009cAsym’] space*. First, a reference volume and its skull-stripped version were generated using a custom methodology of *fMRIPrep*. Several confounding time-series were calculated based on the *preprocessed BOLD*: framewise displacement (FD), DVARS, and three region-wise global signals. FD and DVARS are calculated for each functional run, using their implementations in *Nipype*^51^. The three global signals are extracted within the CSF, the WM, and the whole-brain masks. Additionally, a set of physiological regressors were extracted to allow for component-based noise correction^78^. Principal components are estimated after high-pass filtering the *preprocessed BOLD* time-series (using a discrete cosine filter with 128s cut-off) for the two *CompCor* variants: temporal (tCompCor) and anatomical (aCompCor). tCompCor components are then calculated from the top 5% variable voxels within a mask covering the subcortical regions. This subcortical mask is obtained by heavily eroding the brain mask, which ensures it does not include cortical GM regions. For aCompCor, components are calculated within the intersection of the aforementioned mask and the union of CSF and WM masks calculated in T1w space, after their projection to the native space of each functional run (using the inverse BOLD-to-T1w transformation). Components are also calculated separately within the WM and CSF masks. For each CompCor decomposition, the *k* components with the largest singular values are retained, such that the retained components’ time series are sufficient to explain 50 percent of variance across the nuisance mask (CSF, WM, combined, or temporal). The remaining components are dropped from consideration. The head-motion estimates calculated in the correction step were also placed within the corresponding confounds file. The confound time series derived from head motion estimates and global signals were expanded with the inclusion of temporal derivatives and quadratic terms for each^79^. Frames that exceeded a threshold of 0.5 mm FD or 1.5 standardised DVARS were annotated as motion outliers. All resamplings can be performed with *a single interpolation step* by composing all the pertinent transformations (i.e. head-motion transform matrices, susceptibility distortion correction when available, and co-registrations to anatomical and output spaces). Gridded (volumetric) resamplings were performed using `antsApplyTransforms` (ANTs), configured with Lanczos interpolation to minimize the smoothing effects of other kernels^80^. Non-gridded (surface) resamplings were performed using `mri_vol2surf` (FreeSurfer).

Many internal operations of *fMRIPrep* use *Nilearn* 0.6.1^81^ (RRID:SCR_001362), mostly within the functional processing workflow. For more details of the pipeline, see the section corresponding to workflows in *fMRIPrep*’s documentation.

### Constrained principal component analysis of emotion identification task data

In order to perform constrained principal component analysis (CPCA)^13,20,22^, we began with (*N * T*) × *P* matrix **X**, containing the BOLD time series for *P* = 214 cortical and subcortical parcels (see “Functional data processing using XCP software”) over *T* = 204 image acquisitions, concatenated across *N* = 116 total subjects from both HC and 22q11.2DS cohorts, such that **X** was 23664 × 214. Next, we constructed an FIR basis set **F** that modeled the *r* = 6 image acquisitions following the *v* = 6 task events (unique combinations of correct, incorrect, and non-responses to threatening or non-threatening stimuli^36,37^) for each of the *N* subjects as a binary indicator, plus an intercept term. Initially, **F** is an (*N * T*) × (1 + (*r * v * N*)) matrix (here, 23664 × 4177). Some subjects were either missing task response data or had a non-response rate of > 30%, so rows of BOLD data in **X** and columns of regressors in **F** were removed. Additionally, some subjects did not have data for a particular response (i.e. a subject identified all stimuli correctly), so we could not construct regressors for them. After these two exclusions, **X** was 21199×214 and **F** was 21199 × 3065. To implement this method while preserving the link between each subject’s responses and their BOLD time series, we operated only on the 21199 non-missing elements of the 23664-row matrix while keeping their original positions in place. Thus, for simplicity and consistency with our implementation, we will describe all matrices in their original size. Moving forward to the first step of the CPCA procedure for extracting task-related signals, we fit the regression equation

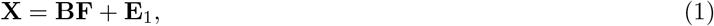

where **B** is a(1 + (*r * v * N*)) × *P* matrix of regression weights from the fitted model, and **E**_1_ is an (*N * T*) × *P* matrix of error terms. The fitted values of Equation 1 contain the variance in **X** that can be explained by the FIR basis set **F** and can be described as

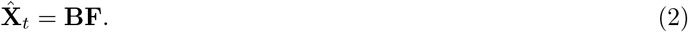

Next, we decompose the task-related variance in 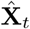 using PCA, following the equation

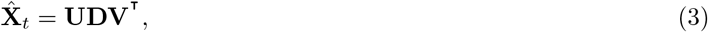

where **U** is an (*N * T*) × (*N * T*) matrix, **D** is an (*N * T*) × *P* diagonal matrix of singular values associated with each component, and **V** is a *P* × *P* matrix whose columns contain orthonormal spatial weights on brain regions for each component. We obtain **Y**, the (*N * T*) × *P* matrix of temporal weights of each component at each modeled BOLD time point, by projecting 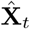 into the space defined by **V**, as described by the equation

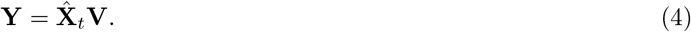

Finally, in order to relate the temporal weights of each PC to the task events, we perform a second regression step by fitting the equation

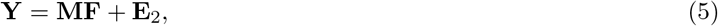

where **M** is a (1 + (*r * v * N*)) × *P* matrix of regression weights from the fitted model, and **E**_2_ is an (*N * T*) × *P* matrix of error terms. **M** contains the estimated temporal response of each of the *P* principal components to each of the *v* task events for each of the *N* subjects.

### Bootstrapping analysis of CPCA components

In order to facilitate the interpretation of these task-evoked modes of brain activity and utilize them to understand potential alterations in brain dynamics in 22q11.2DS, we performed two critical quality control analyses. First, we obtained distributions of each element of the spatial loadings by repeating the entire CPCA procedure using 10,000 bootstrapped samples of the entire dataset. We used these empiric confidence intervals to compute a two-tailed *p*-value for each element of the first 6 spatial loadings, and subsequently thresholded the spatial maps at *p* < 10^−4^ (Fig. 2a for PCs 1-5, and Fig. S1) for the purposes of determining the extent of positive or negative spatial loading in each component not attributable to sampling error.

Second, because we sought to compare temporal expression of group-defined spatial modes of activity, it was important to ensure that group spatial differences did not underlie group temporal differences. To test for this possibility, we performed the CPCA procedure on each group separately and computed the variance explained in bootstrapped samples of each cohort’s BOLD signal by the principal axes of the opposite group, as well as by the group solution. We found that both the group-derived spatial components and the cohort-specific spatial component explained similar amounts of variance in each cohort’s BOLD data, suggesting that aggregating the two groups to identify a common set of axes is an appropriate approach (Fig. S3a). For each component, the dark blue bar and dark red bar are approximately the same height, indicating that over many bootstrapped samples, a similar amount of variance is explained in 22q11.2DS BOLD data by components obtained from the full sample as is explained by components obtained from individuals with 22q11.2DS only. Similarly, the light blue and light green bars are approximately the same height for each component, indicating that over many bootstrapped samples, a similar amount of variance is explained in HC BOLD data by components obtained from the full sample as is explained by components from HCs only. Note that different amounts of variance can be explained by the group solution in each group due to variable temporal expression of the same spatial component, without necessitating that the model is a poor fit for one group or the other. Overall, these results suggest that the group PCA model fits each cohort as well as each cohort’s PCA model fits its own data.

### Multilevel growth model selection procedure

After obtaining the matrix **M** of the temporal responses of each principal component to each stimulus, we next sought to describe the trajectory of PC activity across time. In doing so, we also sought to identify differences in these trajectories between HCs and individuals with 22q11.2DS, and across the different task events (each of the 4 combinations of correct and incorrect responses to threat and non-threat stimuli). First, we discarded the rows of **M** that corresponded to non-response trials, because while it was important to make sure these trials did not go unmodeled in the CPCA procedure, we did not expect non-response trials to reveal consistent patterns of event-related brain activity across subjects. We also discarded column 1 (corresponding to global signal, shown in Fig. S2) and columns 7 through *P* of **M** in order to model only the first 5 principal components, as determined by the scree plot (Fig. S1a). Next, we carried out a model selection procedure to accurately and parsimoniously model the trajectory of each PC’s activation across time and its moderation by 22q11.2DS status and response type, while controlling for age, sex, total brain volume, mean framewise displacement during task scans, and handedness^35^. In each step of the procedure, we used full maximum likelihood estimation to fit a series of multilevel growth models with the `nlme`^82^ package in R, where the dependent variable is always the estimated response of the ith principal component score, contained in the *i*th column of **M**, and the independent variables were determined by the model selection procedure. We specified a two-level model, where repeated measures of activity following each task event were nested within each participant.

In the model selection procedure, we followed the approach taken by a previous study^35^ to sequentially add predictors while ensuring that the increase in model complexity provided a statistically significant improvement in model fit. In each step, a model with fewer parameters was compared to a model with more parameters. In order to select the more complex model as the new “gold standard,” the more complex model had to satisfy 3 criteria:

1. Lower value of the Akaike Information Criteria (AIC)^83^
2. *p* < 0.05 for the log-likelihood ratio test, to assess whether the log-likelihood of the more complex model is greater than the more simple model
3. *p* < 0.05 for the additional coefficients in the more complex model

Using these criteria for determining the superiority of one model over another, we employed the following standardized, partially supervised model selection procedure, annotated with the functions used to implement them in our publicly available code repository (`code/statfxns/lme_msfxns.R`):

1. *Fit base model* (`lme.ms`): Fit a model Φ_*o*_ with fixed effects for age, sex, total brain volume, mean framewise displacement during task scans, handedness, and group membership (22q11.2DS or control), and random intercepts for subject, as defined by the equation:

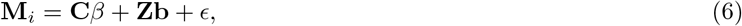

where **M**_*i*_ is the vector of estimated responses for the *i*th principal component, *β* is a vector of optimized regression weights for the *p* fixed effects for all independent variables in the matrix **C, b** is a vector of optimized, subject-specific regression weights that model random effects for time and time polynomials contained in **Z**, and *ϵ* is a vector of normally distributed errors. **b** is normally distributed with 0 mean, and both **C***β* and **Zb** contain intercept terms.
2. *Add fixed effects of time* (`lme.compare`): Compare Φ_*o*_ to *ξ_k_*, where *ξ_k_* contains the parameters of Φ_*o*_ plus a *k*th order polynomial of time as a predictor. Discard all *ξ_k_* that are inferior to Φ_*o*_.

a. *Find the most appropriate model with a significant fixed effect of time* (`lme.selectbest`): Compare the remaining models in *ξ* to one another and set Φ_*t*_ equal to the most superior *ξ_k_*, which contains a polynomial of order *t*.
b. *Add fixed interactions between time, stimulus type, and response type* (`lme.stepdown`): Compare Φ_*t*_ to *ξ_k_*, where *ξ_k_* contains the parameters of Φ_*t*_ plus 3-way interactions between response type, stimulus type, and time polynomials from order *k* to order 0, where the maximum value of *k* is *t*. Set Φ_*t*_ equal to *ξ_k_* with the largest value of *k* for which *ξ_k_* was superior to Φ_*t*_.

i. *Add fixed interactions between time and stimulus type, or between time and response types* (`lme.stepdown`): If there are time polynomial terms without 3-way interactions, compare Φ_*t*_ to *ξ_k_*, where *ξ_k_* contains the parameters of Φ_*t*_ plus 2-way interactions between response type or stimulus type (sequentially) and time polynomials from order *k* to order 0, where the maximum value of *k* is *t*. Set Φ_*t*_ equal to *ξ_k_* with the largest value of *k* for which *ξ_k_* was superior to Φ_*t*_.
c. *Add random effects of time* (`lme.stepup`): Compare Φ_*t*_ to *ξ_k_*, where *ξ_k_* contains the parameters of Φ_*t*_ plus random effects of time from order 1 to order *k*, where the maximum value of *k* is *t*. Set Φ_*t*_ equal to *ξ_k_* with the largest value of *k* for which *ξ_k_* was superior to Φ_*t*_.
d. *Add interactions between 22q11.2DS status and time* (`lme.selectbest`): Compare Φ_*t*_ to *ξ_k_*, where *ξ_k_* contains the parameters of Φ_*t*_ plus interactions between group membership and time polynomials from order 1 to *k*, where the maximum value of *k* is *t*. Set Φ_*t*_ equal to the most superior *ξ_k_*.
e. *Add interactions between 22q11.2DS status, time, stimulus type, and response type* (`lme.stepdown`): Compare Φ_*t*_ to *ξ_k_*, where *ξ_k_* contains the parameters of Φ_*t*_ plus 4-way interactions between response type, stimulus type, group membership, and time polynomials from order *k* to order 0, where the maximum value of *k* is *t*. Set Φ_*t*_ equal to *ξ_k_* with the largest value of *k* for which *ξ_k_* was superior to Φ_*t*_.

i. *Add interactions between 22q11.2DS status, stimulus type, and time, or between 22q11.2DS status, response type, and time* (`lme.stepdown`): If there are time polynomial terms without 4-way interactions, compare Φ_*t*_ to *ξ_k_*, where *ξ_k_* contains the parameters of Φ_*t*_ plus 3-way interactions between response type or stimulus type (sequentially), group membership, and time polynomials from order *k* to order 0, where the maximum value of *k* is *t*. Set Φ_*t*_ equal to *ξ_k_* with the largest value of *k* for which *ξ_k_* was superior to Φ_*t*_.

In the above procedure, we began with covariates in the model in case these confounding factors may have obscured a relationship with time. The `lme.selectbest` function was used instead of the `lme.stepdown` or `lme.stepup` functions when it was possible for a model differing by more than one parameter to outperform Φ_*o*_ or Φ_*t*_. This situation occurs when comparing polynomials of time, where it is possible for a linear time model to show equivalent performance with a 0^th^ order no-time model, but a quadratic or higher order model can outperform the no-time model due to the non-linear relationship with time. The coefficient table of the final model for **M**_*i*_ was used to assess the relationships between 22q11.2DS group membership, time, stimulus type, and response type. These tables are attached as Supplementary Data File 1.

## SUPPLEMENTARY DATA FILES

1. Supplementary Data File 1. Coefficients for non-linear mixed effects models of each of the 6 principal component time courses.

## SUPPLEMENTARY FIGURES

**FIG. S1.**
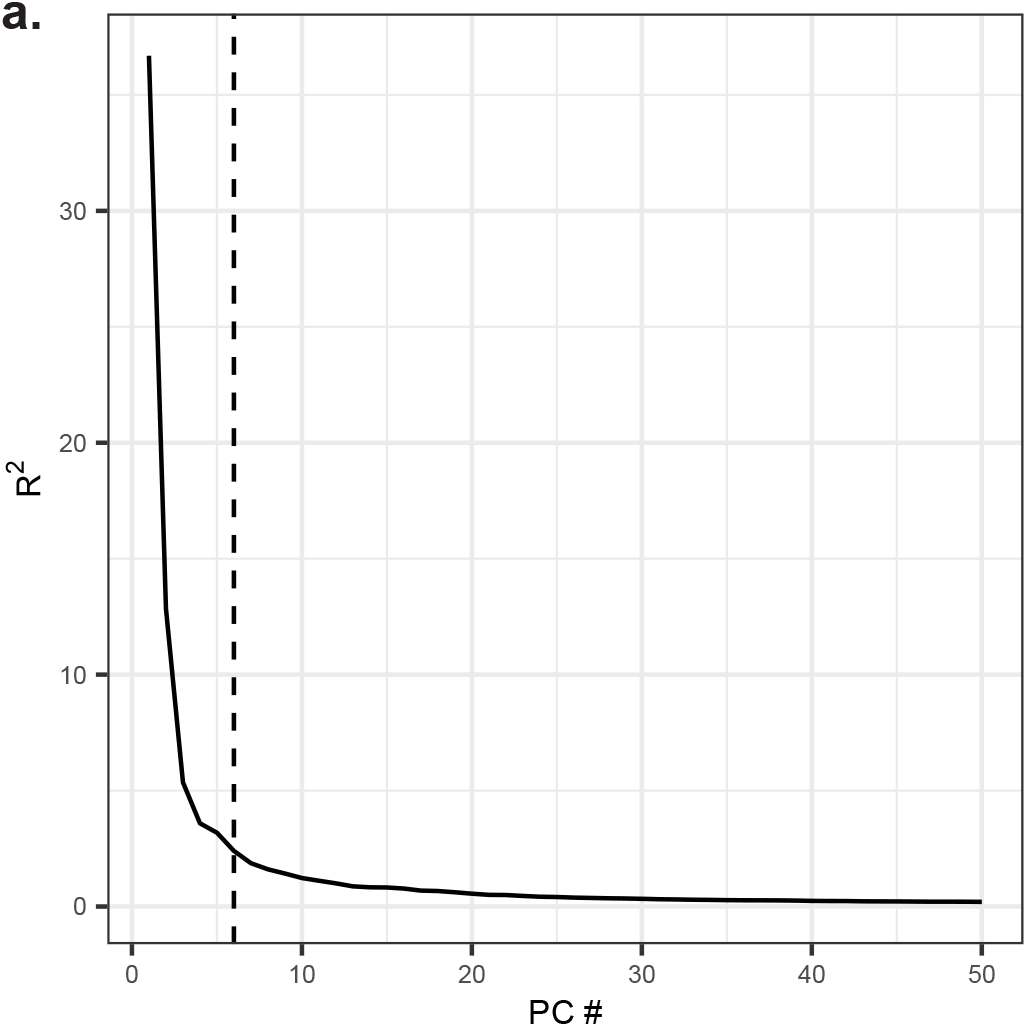
Scree plot of variance explained by PCA step in CPCA procedure. *(a)* Scree plot showing the amount of variance explained (*y*-axis) by each of the first 50 principal components (*x*-axis).

**FIG. S2.**
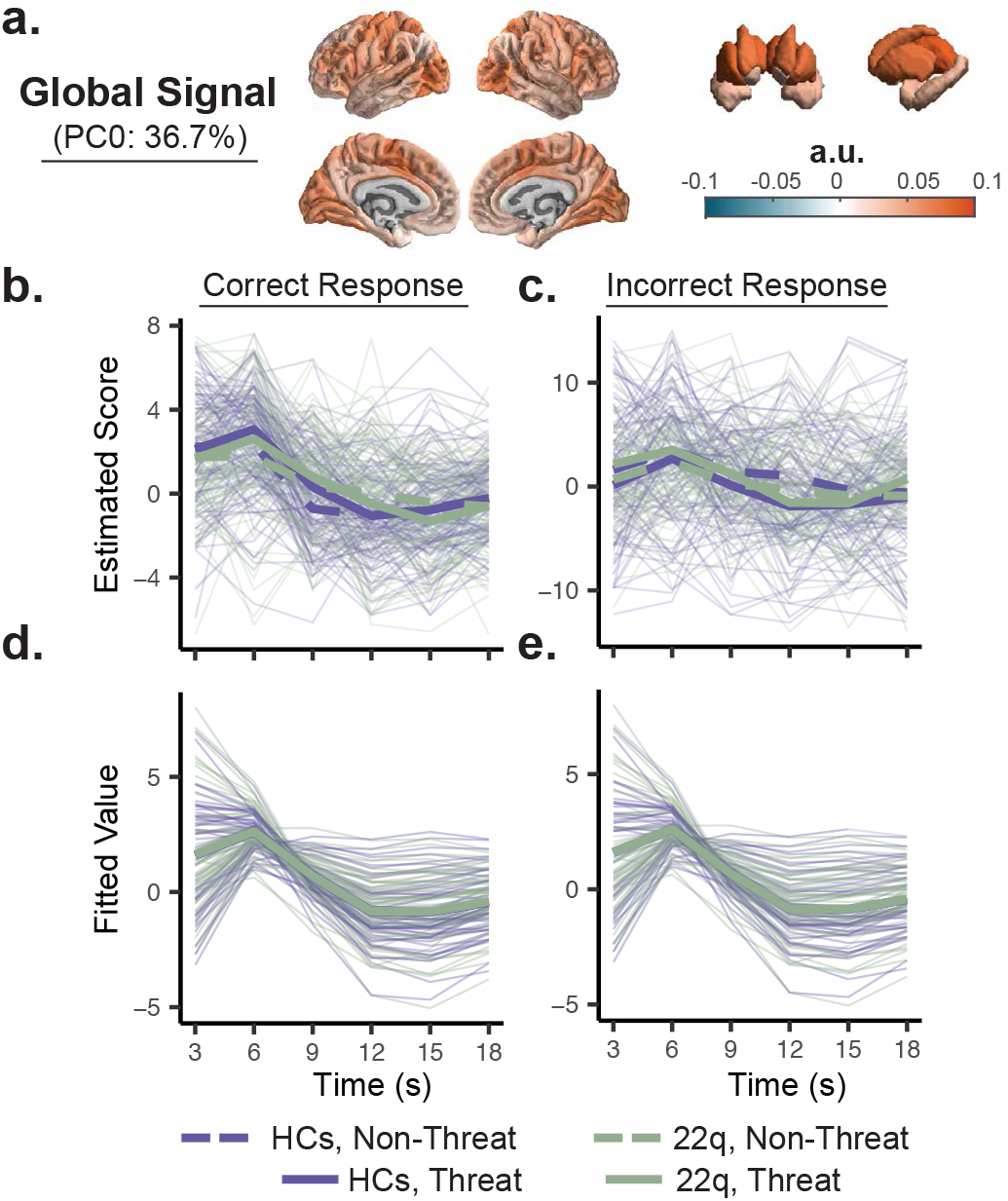
First principal component captures apparent global signal fluctuation. *(a)* Spatial loadings of the first principal component (“PC0”) of task-related variance (Fig. 1b) in emotion identification task BOLD signal reflects apparent global signal fluctuation^38^. Maps were thresholded at *p* < 10^−4^ using bootstrap significance testing^41^ and displayed on surface renderings of cortex and subcortex. *(b, c)* Mean temporal score (*y*-axis) of each task-evoked PC during the time (*x*-axis) period 3-18 seconds after correct (panel *b*) or incorrect (panel *c*) emotion identification of threatening (thick lines) and nonthreatening (dashed lines) faces. The thick lines represent group average values, while the faded lines represent individual subject trajectories. *(d, e)* Multilevel growth models fit to the data in panel *b* (panel *d*) or panel *c* (panel *e*) using the partially supervised model selection procedure described in Methods and Supplementary information. These multilevel models did not contain any coefficients or interaction terms involving 22q11.2DS status.

**FIG. S3.**
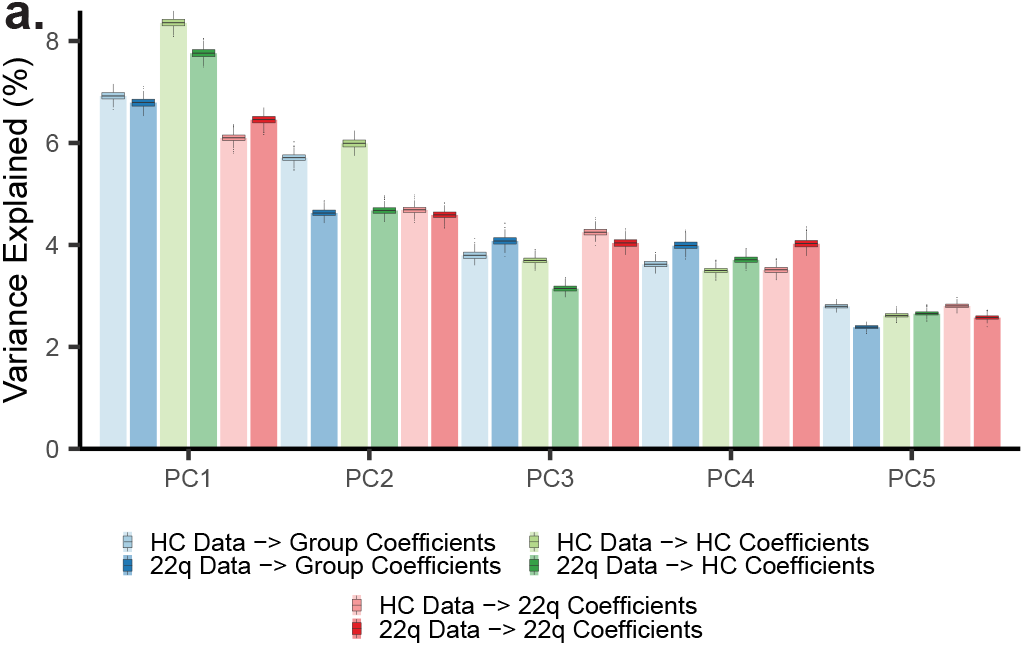
Group principal component analysis solutions capture similar amounts of variance in each cohort. *(a)* Boxplots of explained variance (*y*-axis) by each component (*x*-axis) in either HC or 22q bootstrapped BOLD data by projecting BOLD data into a component space obtained from either HC subjects (“HC Coefficients”), 22q11.2DS subjects (“22q Coefficients”), or all subjects (“Group Coefficients”). This analysis shows that a similar amount of variance in each group is explained by components obtained from either group or the entire sample, suggesting that the group solution is sufficient to explain data from either group.

**FIG. S4.**
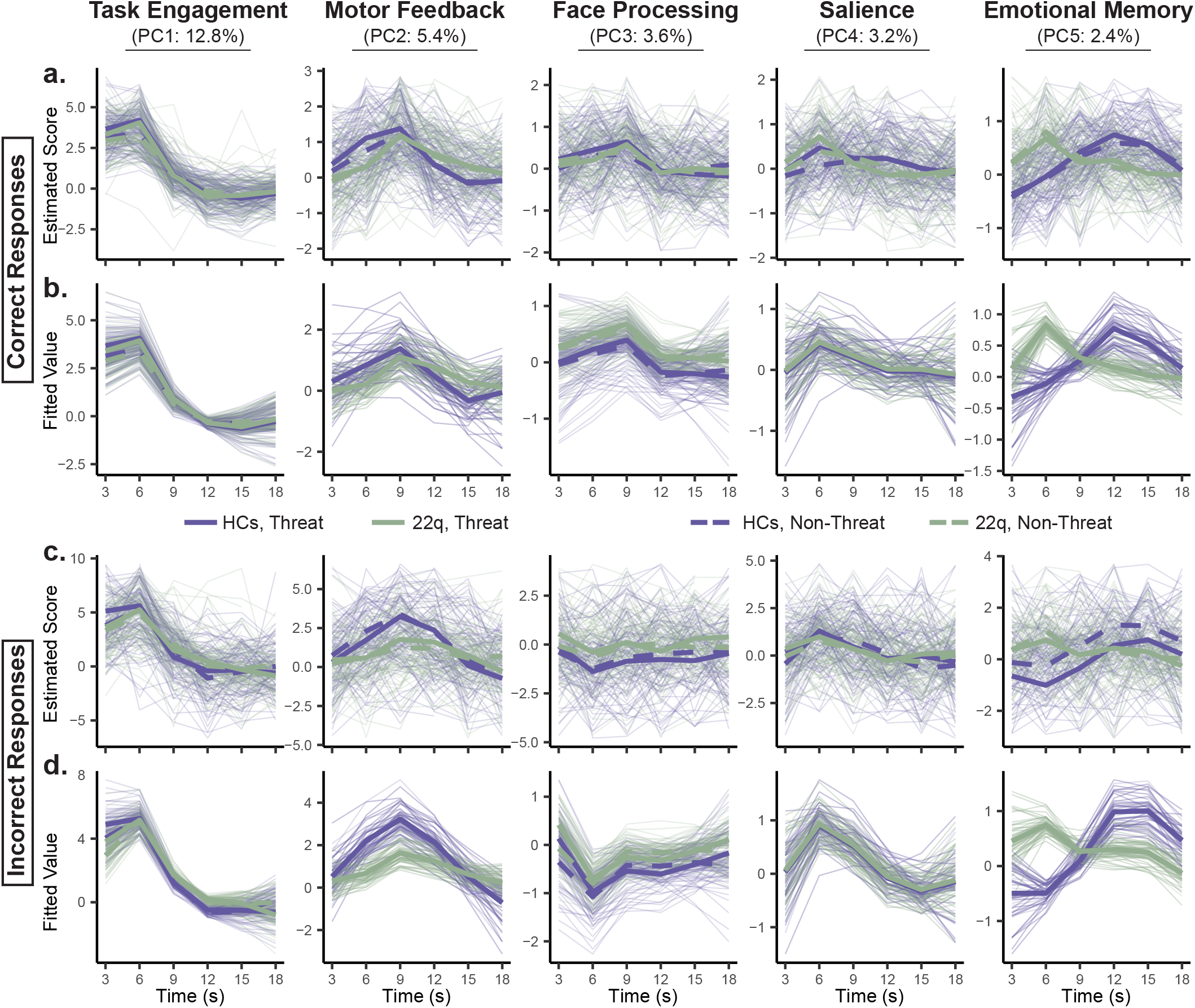
Evoked responses of CPCA components. *(a, c)* Mean temporal score (*y*-axis) of each task-evoked PC during the time (*x*-axis) period 3-18 seconds after correct (panel *a*) or incorrect (panel *c*) emotion identification of threatening (thick lines) and non-threatening (dashed lines) faces. The thick lines represent group average values, while the faded lines represent individual subject trajectories. *(b, d)* Multilevel growth models fit to the data in panel *a* (panel *b*) or panel *c* (panel *d*), reproduced from Figure 2b,c.

**FIG. S5.**
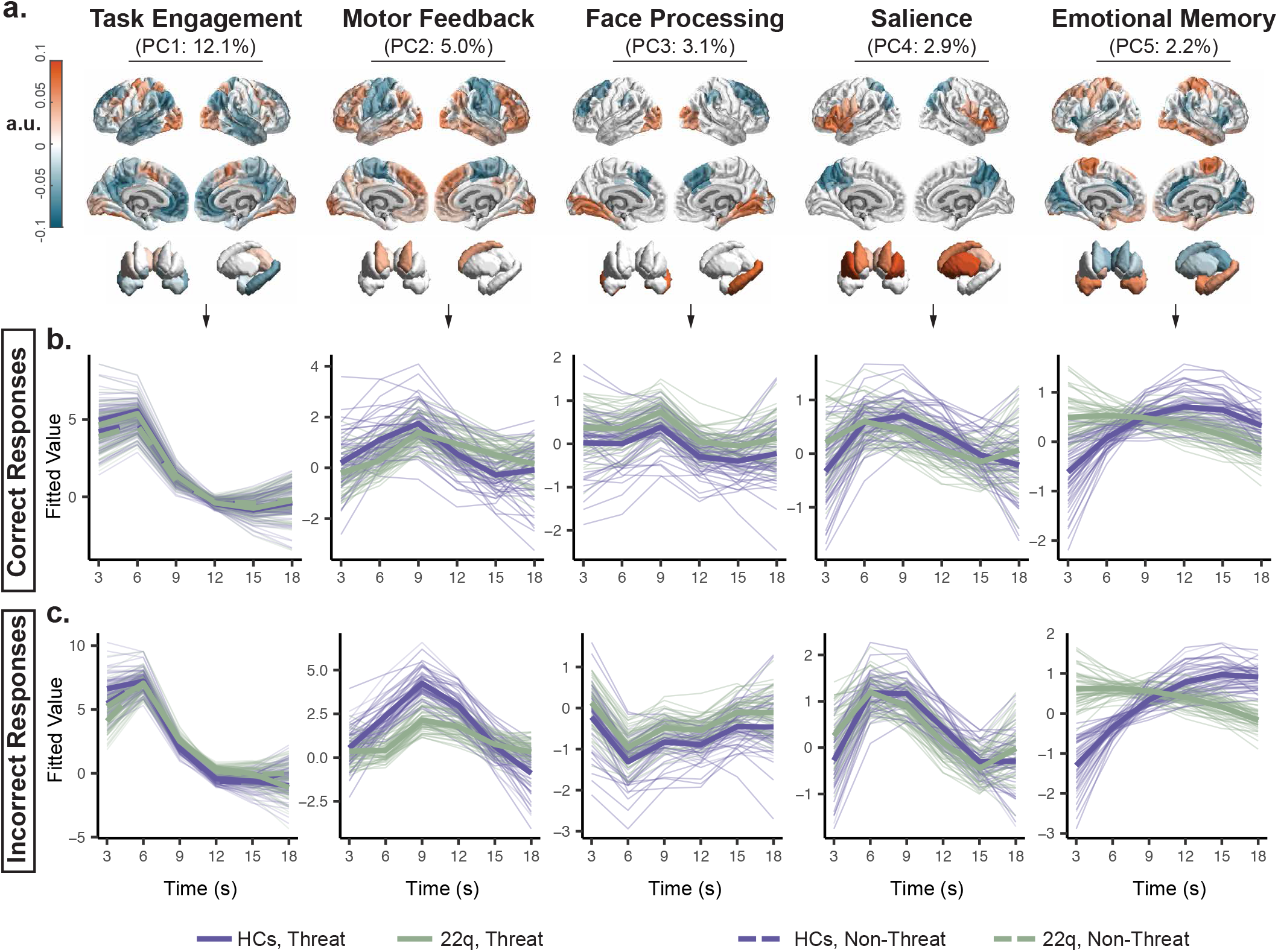
Task-evoked CPCA components using 400 node Schaefer cortical parcellation. This figure is a replication of Fig. 2 using the 400 node Schaefer parcellation^32^ with 14 subcortical nodes defined using the Harvard-Oxford atlas. *(a)* Spatial loadings of the first 5 principal components of task-related variance (Fig. 1b) in emotion identification task BOLD signal thresholded at *p* < 10^−4^ using bootstrap significance testing^41^, shown on surface renderings of cortex and subcortex. Components are named for the cognitive process they putatively reflect. *(b, c)* Multilevel growth models fit to the temporal scores (*y*-axis) of each task-evoked PC during the time (*x*-axis) occurring 3-18 seconds after correct (panel *b*) or incorrect (panel *c*) emotion identification of threatening (thick lines) and non-threatening (dashed lines) faces. We used a model selection procedure (see Methods) to predict each PC’s scores over time from polynomials of time, stimulus type (threat or non-threat), response type (correct or incorrect), 22q status, and interactions between those variables while controlling for age, sex, total brain volume, head motion, and handedness. The best model selected through this process was used to obtain fitted values (*y*-axis) to describe the trajectory of each PC’s score for the prototypical individual in each group (thick, opaque lines) and for each participant (thin, faded lines).

**FIG. S6.**
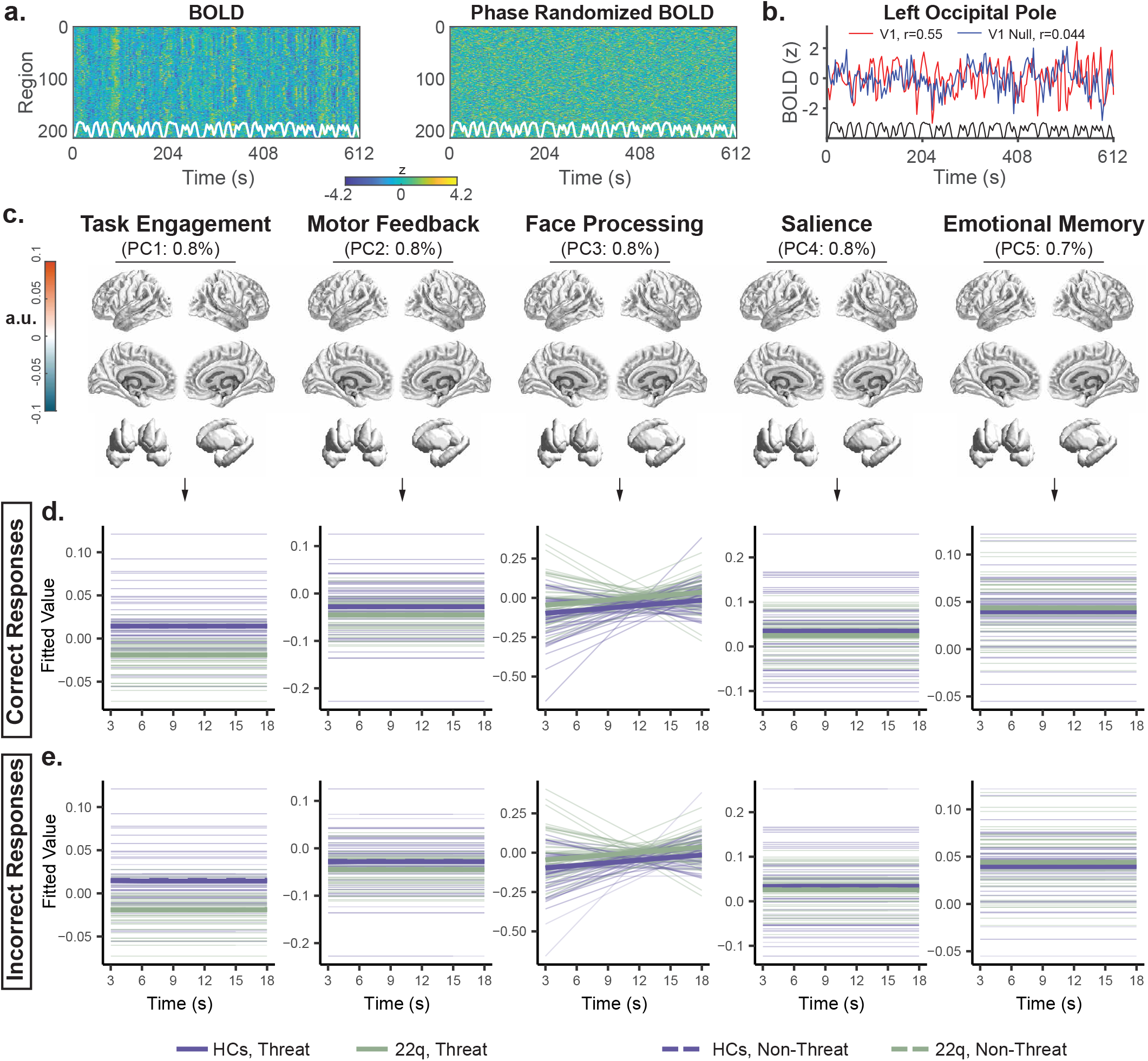
Task-evoked CPCA components with phase randomized BOLD data. This figure is a replication of Fig. 2 with each region’s BOLD time series independently phase randomized within each subject. *(a)* Original BOLD time series (left) next to phase randomized time series (right) for a selected subject. White overlay is the stimulus convolved with canonical hemodynamic response function. *(b)* BOLD time series (red) and phase randomized time series (blue) for left primary visual cortex parcel for selected subject plotted against convolved stimulus (black). Pearson correlations show weaker relationship between stimulus and signal in phase randomized data. *(c)* Spatial loadings of the first 5 principal components of task-related variance (Fig. 1b) in emotion identification task BOLD signal thresholded at *p* < 10^−4^ using bootstrap significance testing^41^, shown on surface renderings of cortex and subcortex. Components are named for the cognitive process they putatively reflect. *(d, e)* Multilevel growth models fit to the temporal scores (*y*-axis) of each task-evoked PC during the time (*x*-axis) occurring 3-18 seconds after correct (panel *d*) or incorrect (panel e) emotion identification of threatening (thick lines) and non-threatening (dashed lines) faces. We used a model selection procedure (see Methods) to predict each PC’s scores over time from polynomials of time, stimulus type (threat or non-threat), response type (correct or incorrect), 22q status, and interactions between those variables while controlling for age, sex, total brain volume, head motion, and handedness. The best model selected through this process was used to obtain fitted values (*y*-axis) to describe the trajectory of each PC’s score for the prototypical individual in each group (thick, opaque lines) and for each participant (thin, faded lines).

